# Estimating axial diffusivity in the NODDI model

**DOI:** 10.1101/2020.10.09.332700

**Authors:** Amy FD Howard, Michiel Cottaar, Mark Drakesmith, Qiuyun Fan, Susie Y Huang, Derek K Jones, Frederik J Lange, Jeroen Mollink, Suryanarayana Umesh Rudrapatna, Qiyuan Tian, Karla L Miller, Saad Jbabdi

## Abstract

To estimate microstructure-related parameters from diffusion MRI data, biophysical models make strong, simplifying assumptions about the underlying tissue. The extent to which many of these assumptions are valid remains an open research question. This study was inspired by the disparity between the estimated intra-axonal axial diffusivity from literature and that typically assumed by the Neurite Orientation Dispersion and Density Imaging (NODDI) model (*d*_║_ = 1.7μm^2^/ms). We first demonstrate how changing the assumed axial diffusivity results in considerably different NODDI parameter estimates. Second, we illustrate the ability to estimate axial diffusivity as a free parameter of the model using high b-value data and an adapted NODDI framework. Using both simulated and *in vivo* data we investigate the impact of fitting to either real-valued or magnitude data, with Gaussian and Rician noise characteristics respectively, and what happens if we get the noise assumptions wrong in this high b-value and thus low SNR regime. Our results from real-valued human data estimate intra-axonal axial diffusivities of ~ 2 – 2.5μm^2^/ms, in line with current literature. Crucially, our results demonstrate the importance of accounting for both a rectified noise floor and/or a signal offset to avoid biased parameter estimates when dealing with low SNR data.

## 1. Introduction

In diffusion MRI, biophysical models aim to relate macroscopic diffusion signals to microscopic, biologically meaningful tissue parameters such as fibre orientation, dispersion or diameter. The primary challenge for biophysical models is being able to describe the complex tissue microstructure in only a handful of parameters that can be estimated reliably from the diffusion signal. Consequently, the model must make strong, simplifying assumptions about the underlying architecture and/or diffusion properties of the tissue. Some parameters are often constrained or set to a single value which is either taken from the literature or decided by some other means (e.g. by fitting the model multiple times with different values for some assumed parameter, and taking the result which maximises the quality of the fit across voxels or specimens[1]). Though these assumptions make the estimation of otherwise degenerate parameters tractable, if the modelling constraints are inaccurate, the estimation of the remaining model parameters will be biased [2, 3].

Neurite Orientation Dispersion and Density Imaging (NODDI) [4] is a commonly used biophysical model in diffusion MRI which, due to its clinically feasible scan times requirements, has been reported in numerous studies of patient populations [5, 6, 7, 8, 9]. NODDI is a variant of the white matter ‘standard model’ [10] in which specific assumptions about the tissue diffusivity allow for voxelwise estimation of fibre dispersion and neurite ‘density’ or signal fraction. One assumption is that the intra-axonal axial diffusivity, i.e. the diffusion of water molecules inside the axon as they travel along the primary axis or orientation, is a fixed, global value known *a priori* and typically set to 1.7 μm^2^/ms. However, there now exists a body of work in which the intra-axonal axial diffusivity is generally estimated to be higher, typically in the range of ~ 2 — 2.5μm^2^/ms [11, 12, 2, 13, 14, 15, 16, 17, 18, 19]. Notably, these studies use different methods to achieve compartment specific selectivity (including high b-value data [16, 20], gadolinium injection [15] and planar filtering [17]), as well as different microstructure models and/or parameter constraints to estimate axial diffusivity.

Inspired by the difference between the reported and assumed intra-axonal axial diffusivity, this study first aims to explore how changing the predefined axial diffusivity in the NODDI model affected the remaining estimated parameters. A second aim of this study was to investigate whether, by utilising high b-value data, we could simultaneously estimate axial diffusivity within the NODDI framework. Here it is useful to estimate axial diffusivity within the NODDI framework (rather than e.g, from the voxels with highest fractional anisotropy [21]) as it facilitates estimation of both axial diffusivity and fibre dispersion on a voxelwise basis. Were the fibres instead assumed to be coherently aligned, the axial diffusivity estimates would be biased [3, 17]. Crucial to our second aim was the use of ultra-high b-value data, where it can be assumed that the higher-diffusivity extra-axonal water is eliminated such that only the intra-axonal compartment contributes signal [22, 23, 16, 24, 20, 25, 26], thus overcoming known degeneracies between diffusion characteristics of the intra- and extra-axonal space [3]. However, high b-value data also posed several challenges. In particular, we found that in this low SNR regime, the model had to account for both the rectified noise floor and/or a signal offset to avoid parameter degeneracy and bias. We demonstrate how, with appropriate modifications, the NODDI framework can be applied to both magnitude and real-valued data to estimate axial diffusivities in line with previous literature.

## 2. Methods

This paper is organised as follows. First we demonstrate how changing the assumed intra-axonal axial diffusivity affects the output of the ‘standard NODDI model’ (described below), highlighting the importance of the discrepancy between the assumed axial diffusivity of NODDI and many estimates of axial diffusivity found in literature. We then investigate how a ‘modified NODDI model’ can be used to estimate both the axial diffusivity and ODI concurrently from high b-value data. Specifically, this modified NODDI model (i) considers the intra-axonal compartment only and (ii) explicitly accounts for Rician noise floor and a signal offset. Finally, the modified model was applied to *in vivo,* human data, where we explore how noise can bias model estimates in this high b-value and thus low SNR regime.

### 2.1. NODDI output sensitivity to the assumed axial diffusivity

To evaluate how the NODDI output changed with respect to the assumed axial diffusivity, the standard NODDI model [4] was applied to the diffusion data with *d*_║_ = 1.7, 2.3 or 3 μm^2^/ms. Briefly, the NODDI model consists of a Watson-like fibre orientation distribution which is convolved with three compartments that are typically associated with the CSF, extra-axonal and intra-axonal space. The first compartment has isotropic, free diffusion, the second compartment describes tensor-like diffusion, and the third compartment describes stick-like diffusion. The model fitting involves 5 parameters being estimated (the intra-axonal signal fraction, the isotropic signal fraction, the fibre orientation and orientation dispersion index, [*f_in_, f_iso_,θ,ϕ,ODI*]) whilst assuming some fixed, global axial diffusivity which may differ from *d*_║_ = 1.7μm^2^/ms, that the radial diffusivity of the tensor-like compartment is given by the tortuosity model *d*_⊥_ = *d*_║_ (1 — *f_in_*) and that the diffusivity of the isotropic compartment is that of free diffusion *d_iso_* = 1.7 μm^2^/ms.

Here we utilised preprocessed T1-weighted and diffusion-weighted data for the first 10 subjects of the WU-Minn Human Connectom Project (HCP); for details of the acquisition protocol and preprocessing pipeline, please see [27, 28, 29]. Briefly, the diffusion-weighted data included 90 gradient directions each at b-values of *b* = 1, 2 and 3ms/μm^2^ and 18 interspersed volumes with negligible diffusion weighting. The distortion corrected (“pre-processed”) *b* 0 ms/μm^2^ data were linearly registered to each subject’s T1-weighted structural scan (FLIRT [30, 31]), and the T1 non-linearly registered to the MNI standard space (FNIRT [32, 33]). The NODDI fitting [4] was performed in subject space using the cuDIMOT framework [34] for GPU acceleration with Rician noise modelling.

### 2.2. Modified NODDI for high b-value data

The NODDI model was modified for high b-value data where we assume the entirety of the diffusion signal can be attributed to the intra-axonal compartment [22, 23, 16, 24, 20, 25, 26]. Here, the diffusion signal along gradient direction ***g*** is given by the convolution of the fibre orientation distribution, which was assumed to be a Watson distribution [35, 4], and a fibre response function for stick-like fibres with Gaussian axial diffusion and no radial diffusion:

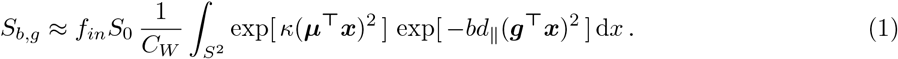

The integrand is over ***x*** ∈ *S*^2^; *f_in_* being the signal fraction of the intra-axonal compartment; ***μ***(*θ,ϕ*), the direction of the fibre; *d*_║_, the intra-axonal axial diffusivity; ***x***, a unit vector on the sphere *S*^2^; *b*, the b-value and *S*_0_, the non-diffusion weighted signal. *C_W_* is the normalising constant, where,

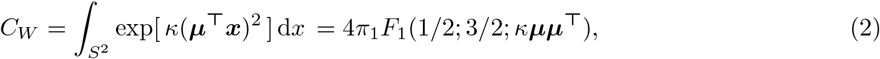

and _1_*F*_1_(*α; β*; ***X***) is the confluent hypergeometric function of the first kind with a matrix argument ***X***. The dispersion parameter *κ* is typically rewritten in terms of the orientation dispersion index, *ODI* = 2/*π* arctan(1/*κ*) [4] which ranges from 0 to 1, representing perfectly aligned and isotropic fibre distributions respectively.

The integral in Equation 1 can also be written in terms of the confluent hypergeometric function of the first kind with matrix argument 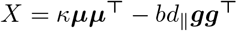. Consequently, Equation 1 becomes [36]:

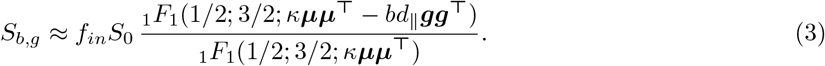

By taking the ratio of 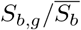 (rather than *S_b,g_*/*S*_0_) we can derive an analytic solution where the model is independent of both *f_in_* and *S*_0_, and described by only four parameters 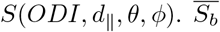 represents the powder averaged signal, i.e. the average signal across all diffusion gradients for a given b-value, where at high b-value, for stick-like diffusion convolved with an arbitrary fibre orientation distribution [13],

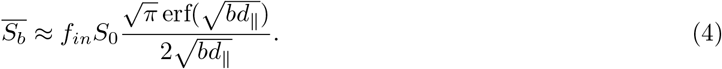

Dividing *S_b,g_* (Equation 3) by 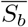 (Equation 4), the diffusion signal has an analytic form given by:

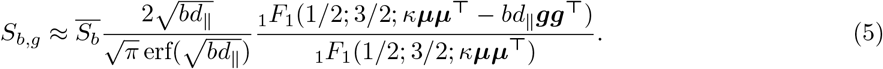

Here the model depends on only two free parameters, *d*_║_ and *ODI* (or *κ*). During fitting, 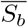 was calculated for each shell in turn.

In our initial investigations we aimed to simultaneously estimate the three parameters *d*_║_, *κ* and *F_in_* = *f_in_S*_0_ on a voxelwise basis according to Equation 3. However, upon closer examination, the parameters were found to be degenerate, leading to an overestimation of *d*_║_. Here we used simulated data (shown later) to investigate the model parameter degeneracy and bias in relation to the noise characteristics of the data. Crucially, we found that the parameter degeneracies and bias could be overcome if we fitted to high SNR data, and explicitly model both a signal offset and the Rician noise floor. The presence of a signal offset in the *in vivo* data was supported by the presence of non-zero signal in the ventricles at high b-value. Consequently, we accounted for a signal offset i.e. some background signal that was independent of diffusion weighting, where

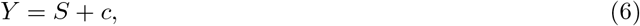

*Y* being the data, *S* the diffusion signal and *c* the offset, which is sometimes referred to as a dot compartment. Then in magnitude data, we also accounted for the rectified Rician noise floor using Koay’s inversion technique [37]:

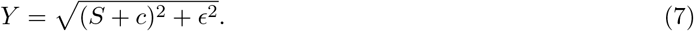

*ϵ* being the Rician scaling parameter, which is equivalent to the standard deviation of Gaussian noise for complex data. Typically, *ϵ* is estimated *a priori* from noise estimation methods [38]. However, as shown below, *a priori* estimation might not be ideal as even a small misestimation of e could lead to large parameter biases. An alternative way of circumventing Rician noise floor effects is to consider real-valued data with Gaussian noise characteristics [39]. Using complex data, the signal phase is first removed from each voxel after which the real component of the signal is extracted [39]. In this study, the model was fitted to both real valued data (where *c* was estimated) and magnitude data (where both *c* and *ϵ* were estimated) from the same subjects. As even a small misestimation of *c* or *ϵ* can lead to a large parameter bias, both were estimated as parameters of the model that were fitted voxelwise during model optimisation.

#### 2.2.1. The final model & optimisation

Combining Equation 5 with either Equation 6 (real-valued data) or 7 (magnitude data) and assuming a known fibre orientation *μ* = *V*_1_ (the primary eigenvector of the diffusion tensor [21]), the final model was dependent on only three or four parameters: the orientation dispersion index *ODI*, the axial diffusivity *d*_║_, the signal offset *c*, and, in magnitude data only, the noise floor parameter *ϵ*. Figure 1 shows a graphical depiction of the model.

**Figure 1:**
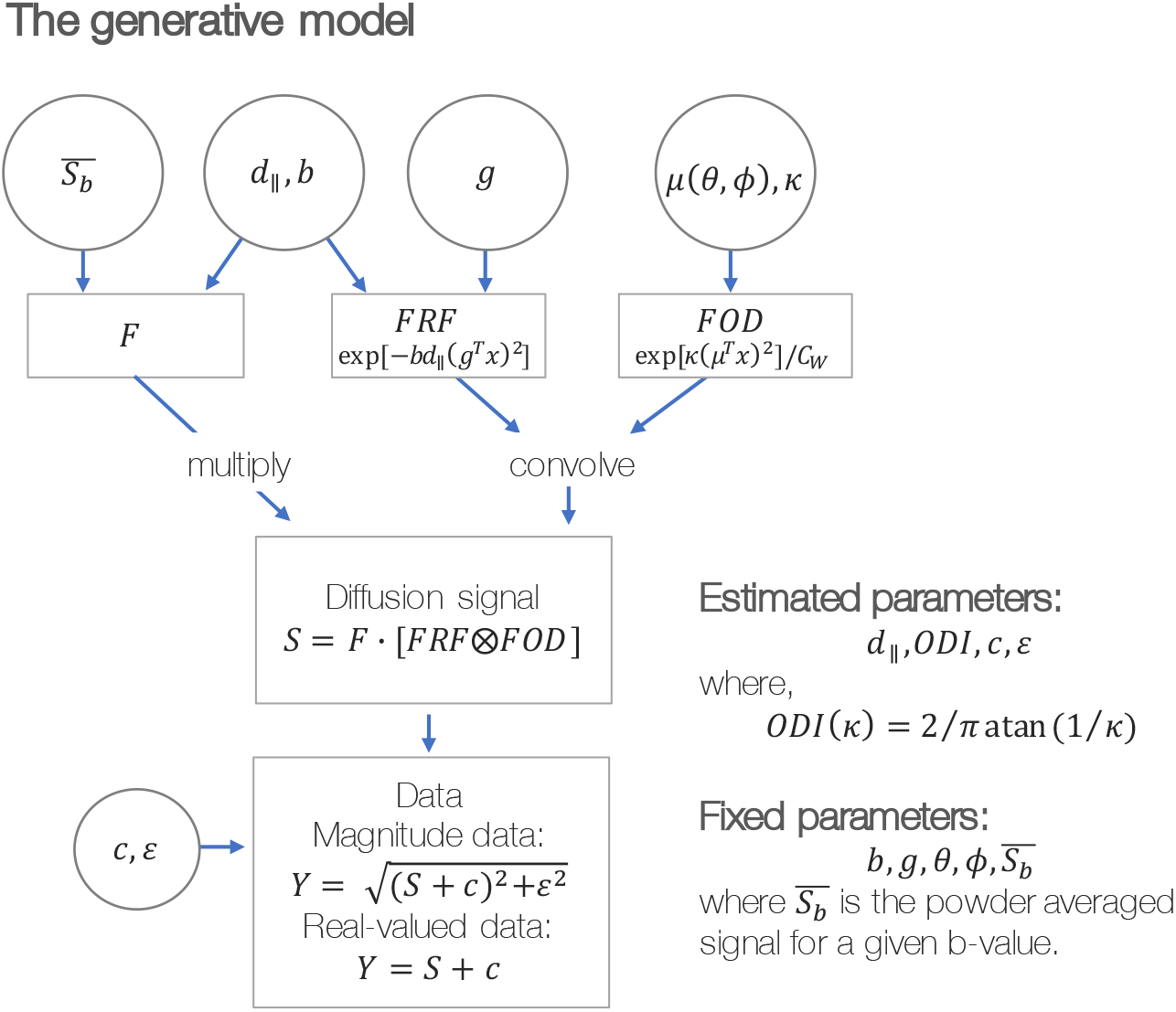
The modified NODDI model for co-estimation axial diffusivity and orientation dispersion in high b-value data. The fibre response function (*FRF*) and fibre orientation distribution *(FOD)* can be first convolved and then multiplied by the non-attenuated diffusion signal associated with the intra-axonal compartment (*F* = *f_in_* · *S*_0_) to produce the diffusion signal *S*. A signal offset *c*, and (in magnitude data only) rectified noise floor *ϵ*, were then added to the signal to produce the real or magnitude data, *Y* as required. Here we use the powder averaged signal 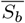 to avoid estimating *F* as a parameter of the model. See the main text for full definition of parameters.

During optimisation, the parameters were bounded such that *ODI* ∈ [0,1] and *d*_║_ ∈ [0, 4] μm^2^/ms where the diffusivity in free water at 37°C is ~ 3 – 3.1 μm^2^/ms. Due to the observed possible degeneracy between the two noise parameters *c* and *ϵ*, when both were estimated in magnitude data, they were constrained to be 50 — 150% of c and e as estimated from high b-value data in the ventricles (a Rician distribution was fitted to *b* = 17.8ms/μm data, where *c* = the non-centrality parameter and *ϵ* = the scale parameter). When only one noise parameter was estimated (in real-valued data or simulations with *c* = 0), the noise parameter was constrained such that *c* ∈ [0,0.5 · So] or *ϵ* ∈ [0, 0.5 · *S*_0_]. The powder averaged signal 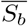 was calculated for each shell in turn, where the data *Y* were for first corrected for any signal offset *c* or noise floor e according to Equation 6 or 7 (i.e. *Y* was converted to *S* prior to signal averaging). The model was optimised using the Metropolis Hastings (MH) algorithm, which afforded estimation of each parameter’s posterior distribution. The initial parameters for MH were found by grid search.

Our investigations using simulated data demonstrated how higher SNR leads to more precise estimates of *ODI* and *d*_║_. Consequently, for *in vivo* data, the model was fitted to the concatenated signal across many (*N*) voxels to boost SNR. Here, the signal from each voxel was first rotated such that the primary eigenvector of the diffusion tensor was aligned and the secondary eigenvector was randomly orientated, since the Watson distribution describes symmetric dispersion about the primary fibre orientation. As NODDI assumes a single-fibre population per voxel, we fitted to high FA voxels from the corpus callosum which, to enforce some data consistency, were selected to have similar *S*_0_ 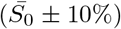. Instead of concatenating the signal, we could have averaged the signal across voxels (as we did using simulated data below). However, this would require 1) the diffusion signal from each voxel to first be rotated so that the primary fibre orientations align, and 2) the rotated signal to be resampled along consistent gradient orientations *g*. Consequently, signal concatenation was deemed preferable to signal averaging across voxels, as our results would not be biased by interpolation effects which can introduce smoothing and effect noise properties in non-trivial ways.

### 2.3. Simulated data

Data were simulated with a known ground truth to investigate the precision and accuracy of the parameter estimates. The *S*_0_, b-values and gradient directions were chosen to mimic the *in vivo* data below. The ground truth parameter values were *d*_║_ = 2.2, *ODI* = 0.03, *f_in_* = 0.6 and *c* =10 unless otherwise stated. The SNR of a single voxel was defined as SNR_*vox*_ = *S*_0_/*σ* = 16.5, which is similar to the lower bound of the SNR in the *in vivo* data used later in this study, where *σ* represents the standard deviation of Gaussian noise in the complex data. As in *in vivo* data, the model was fitted to data from N simulated voxels. The SNR across many voxels was then approximated as 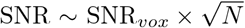.

Note, for *in vivo* data we fitted the model to the concatenated signal across voxels, whilst for simulated data we fitted to the averaged signal. In simulated data, we fixed the fibre orientation and could thus compute the average signal across voxels without interpolation. Furthermore, by averaging the signal, we fitted to fewer gradient directions and minimised computation time.

### 2.4. In vivo human data

The modified NODDI model was applied to pre-existing data from 6 healthy participants where both magnitude and real-valued diffusion images were reconstructed from the complex MR data.

Diffusion MRI data were previously acquired, reconstructed and pre-processed according to Tian *et al.* [40] and Fan *et al.* [39]. Briefly, complex diffusion data were acquired on the 3T MGH Connectom scanner (Siemens Healthcare, Erlangen, Germany) using gradients up to 300mT/m [41, 42] and a pulsed gradient, spin echo EPI sequence [43]: *TR/TE* = 4000/77 ms, 2mm isotropic resolution, 17 b-values from 0 to 17.8ms/μm, two different diffusion times *δ*/Δ = 8/19 ms and *δ*/Δ = 8/49 ms. The complex images were first corrected for background phase contamination after which the real component of the diffusion images was extracted and the imaginary part discarded [39]. The magnitude images were also extracted. The data were pre-processed using tools from FSL and bespoke code. The real-valued and magnitude data were separately corrected for gradient nonlinearities as well as susceptibility and eddy current distortions [44, 45, 46, 47]. Here we fitted the modified NODDI model to data from 6 subjects with Δ = 49 ms, *b* = 6.75, 9.85 and 13.5ms/μm^2^ and 64 gradient directions per shell. The *b* = 0ms/μm^2^ (30 repeats) and *b* = 0.95ms/μm^2^ images with 32 gradient directions were used for fitting the diffusion tensor model only [21].

T1-weighted structural images were also acquired and here used for white matter segmentation [48] and to register a corpus callosum mask from MNI standard space to subject space [32, 33].

Figure 2 shows example preprocessed diffusion images. In Figure 2 right we see how the images retain good signal at very high b-values. Figures 2 left shows the distribution of high b-value (*b* = 17.8ms/μm^2^) signal from voxels in the ventricles for a single subject. The noise from the magnitude images follows a Rician distribution, whilst that from real-valued data is Gaussian distributed as expected. The Rician noncentrality parameter and Gaussian mean are both non-zero and indicative of a positive signal offset. The data SNR, here taken to be *S*_0_/*σ*, where *σ* was taken to be the Guassian standard deviation or Rician scaling parameter, was found to be *SNR* ∈ [17, 30] with a median *SNR* of 21.

**Figure 2:**
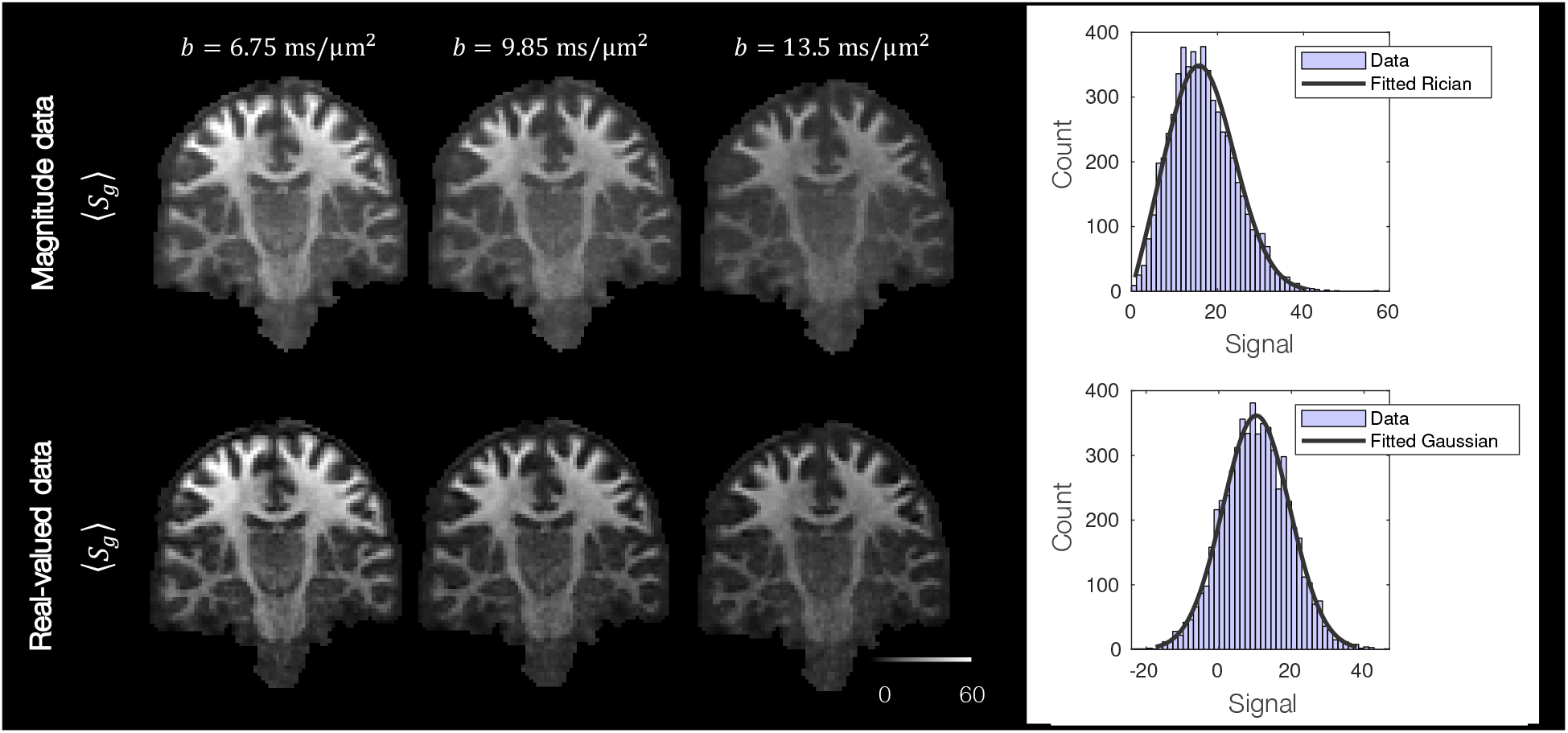
Left: Example magnitude and real-valued diffusion images from a single subject. Here we show the mean signal across all gradient directions. We observe a considerable rectified noise floor in the magnitude images, which appears greatly reduced in the real valued data. Right: The distribution of signal from voxels in the ventricles at *b* = 17.8ms/μm^2^, where we assume the signal to be purely noise. In both cases, the signal is not zero-mean: the magnitude data follows a Rician distribution with the non-centrality parameter *v* = 13.7, and *σ* = 8.4; the real-valued data has Gaussian noise with a positive offset, *μ* = 10.4, *σ* = 9.2. Consequently a signal offset and, in the case of magnitude data, rectified noise floor were estimated as parameters of the model.

## 3. Results

### 3.1. Changes in the NODDI output for different axial diffusivity

Figures 3 and 4 show how the parameters of NODDI [4] change when the assumed axial diffusivity is set to *d*_║_ = 1.7 μm^2^/ms – as is typically assumed – or 2.3 μm^2^/ms, or 3 μm^2^/ms, the latter being the approximate diffusivity of free water.

**Figure 3:**
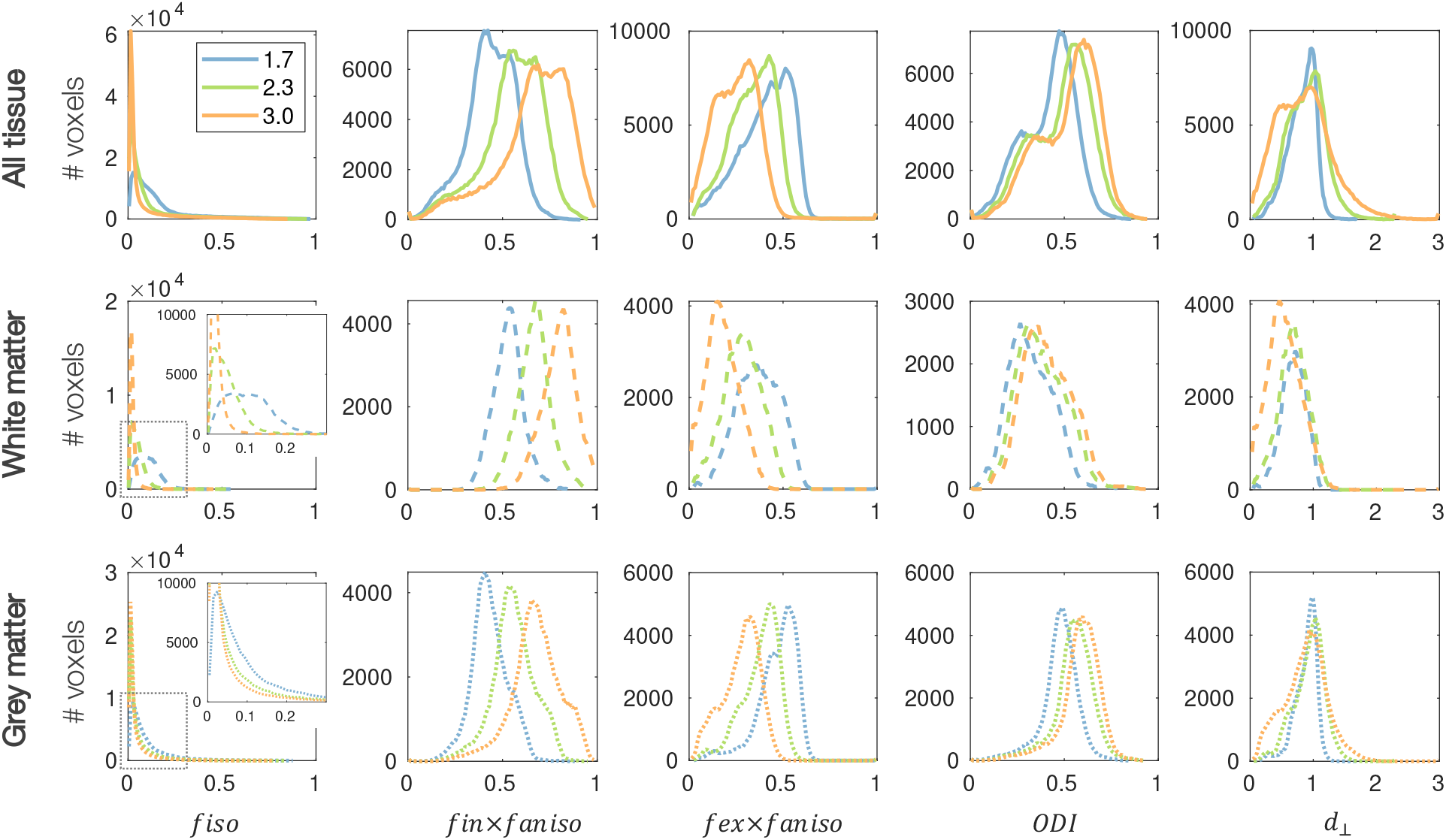
The dependence of NODDI parameters on the assumed axial diffusivity. The NODDI model [4] was fitted to 10 subjects from the HCP dataset with various assumed axial diffusivities *d*_║_ = 1.7, 2.3, 3 μm^2^/ms. The parameter maps were co-registered after which the average map across subjects was calculated. The distribution of these parameters is shown for all voxels in the brain (top) and for the white matter (middle, dashed) and grey matter (bottom, dotted) separately. *f_ansio_* describes the signal fraction of the anisotropic compartment, where *f_ansio_* = 1 – *f_iso_*. *d*_⊥_ is in units of μm^2^/ms.

**Figure 4:**
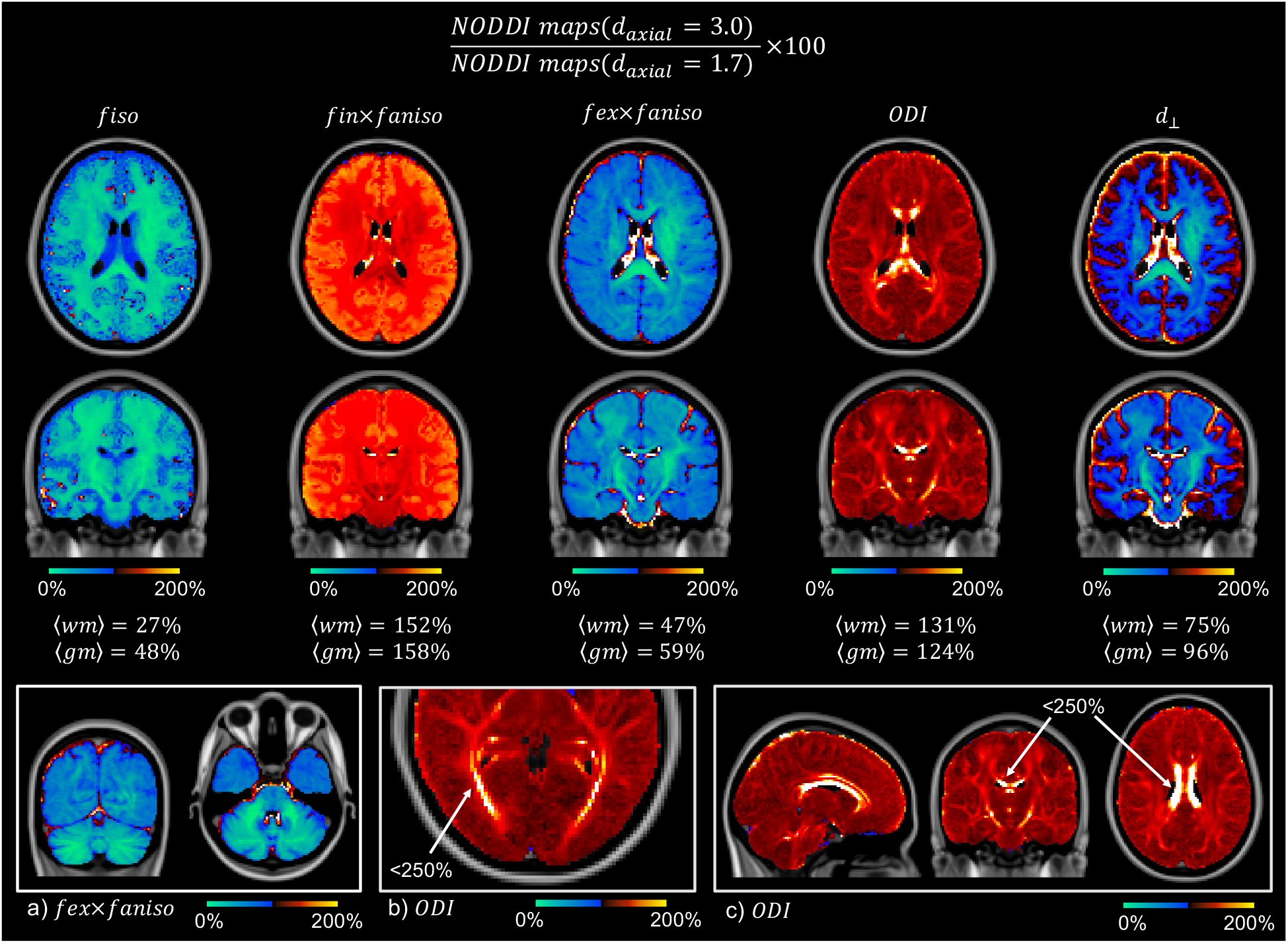
Estimated NODDI parameter maps with axial diffusivity *d*_║_ = 3 μm^2^/ms as a percentage of equivalent maps for *d*_║_ = 1.7 μm^2^/ms, as is typically assumed. The blue shows where parameters decrease, and the red where they increase, as we change *d*_║_ from 1.7 to 3 μm^2^. The signal fraction associated with isotropic diffusivity and the extra-axonal compartment is substantially reduced, as is the radial diffusivity. The intra-axonal signal fraction is largely increased as is the ODI, though to a broadly lesser extent. a) The extra-axonal signal fraction is reduced substantially in the cerebellum. b,c) The ODI increased to < 250% in areas of the optic radiation and corpus callosum.

Figure 3 shows an increase in the gross signal fraction associated with the intra-axonal compartment (*f_in_* × *f_aniso_*) and a decrease of the extra-axonal compartment (*f_ex_* × *f_aniso_*) when the assumed axial diffusivity is increased from 1.7 to 3 μm^2^/ms. Interestingly, when *d*_║_ = 3 μm^2^/ms, both the extra-axonal signal fraction and radial diffusivity (*d*_⊥_) show two distinct distributions associated with the white and grey matter. As *d*_║_ is increased, the ODI is seen to increase in both the grey and white matter, and the signal fraction associated with isotropic diffusion, *f_iso_* is reduced close to zero across most of the brain and particularly in the white matter where we would not typically expect to find isotropic, free diffusion.

In Figure 4 we see the NODDI parameter maps with axial diffusivity *d*_║_ = 3 μm^2^/ms as a percentage of equivalent maps for *d*_║_ = 1.7 μm^2^/ms. Since *d*_║_ = 3 μm^2^/ms represents the upper bound of water diffusion *in vivo*, this comparison represents the maximum difference we may obtain when increasing the assumed axial diffusivity from *d*_║_ =1.7 μm^2^/ms. When *d*_║_ = 3 μm^2^/ms, the isotropic signal fraction and extra-axonal signal fraction decrease on average to 30 — 50% and 50 — 60% respectively of their value when *d*_║_ =1.7 μm^2^/ms. Concurrently, the signal fraction associated with the intra-axonal compartment and the ODI increase on average to ~ 150% and ~ 120%. Here we do not see a global, step change, but rather one which varies across the tissue. In particular, the ODI is substantially increased in the corticospinal tract as well as by 200 — 250% in areas of the optic radiation and corpus callosum. We see a large ODI change across much of the the corpus callosum, though not at the midline along the left-right axis, a known region of increased fibre dispersion.

### 3.2. Modified NODDI model for high b-value data: simulated data

A second aim of this study was to investigate whether, by applying NODDI to high b-value data, it was possible to also estimate axial diffusivity. Crucially, at high b-value, signal contributions from extra-axonal water are assumed negligible and thus the observed signal can be attributed to the intra-axonal compartment, here described by diffusion along dispersed sticks. To examine the accuracy and precision with which this high b-value ‘modified’ model could simultaneously estimate axial diffusivity and ODI, we began by examining simulated data, with known ground truth parameters.

#### 3.2.1. Parameter distributions and model fits

To investigate degeneracy, Figure 5 shows the estimated parameter distributions (left) and model fit (right) for high SNR data (*N* = 100, SNR ~ 165). The model was fitted to data simulated with both a Gaussian and Rician noise model, to mimic real-valued and magnitude diffusion data. In both datasets, the model appeared to fit the data well, where the residuals between the data and the predicted signal were small.

**Figure 5:**
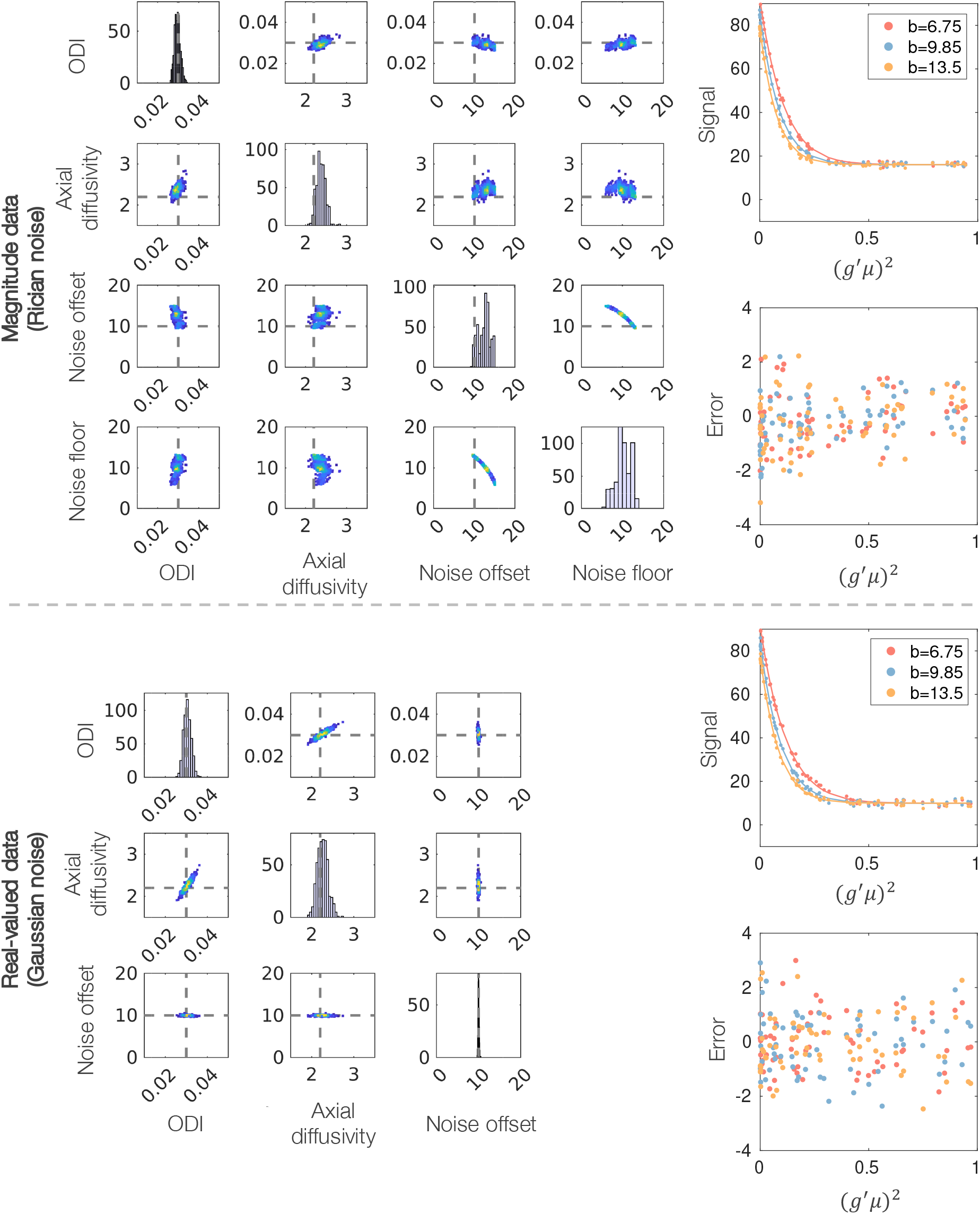
Left: Parameter distributions output from MCMC. Each data point in the density plots represents a combination of parameters that fits the signal equally well. Grey dashed lines represent ground truth values. Right: Model fit (line) to simulated data (dots) and the associated error (prediction-data). *g* is the gradient direction, *μ* the fibre orientation and *b* the b-value in ms/μm^2^. The diffusivities are given in units of μm^2^/ms. The signal is averaged over 100 voxels, each with SNR=16.5. Note, as the noise floor parameter is an approximation, we do not plot its ‘ground truth’ value.

Upon inspection (Figure 2), the *in vivo* data used in this study appeared to contain a signal offset (i.e the complex noise was not zero-mean), that was subsequently included as a parameter of the model (*c*). Figure 5 shows how, in real-valued simulated data, the signal offset can be estimated with ease: it is not correlated to the other model parameters and does not appear to affect their estimation. In magnitude data, the signal offset is highly correlated with the noise floor parameter e, causing difficulties in parameter fitting where multiple minima may exist. In comparison, for data with Gaussian noise (meaning that *ϵ* = 0) the parameter distributions are approximately Gaussian and close to the ground truth values, demonstrating that axial diffusivity and ODI can be estimated reliably and without degeneracy from real-valued data. Supplementary Figure 1 shows similar plots for magnitude data without a signal offset (*c* = 0) where parameter estimation is again improved.

#### 3.2.2. Parameter estimation as a function of SNR

Figure 5 examines parameter estimation in relatively high SNR data. As most *in vivo* data have an *SNR* < 165, Figure 6 shows how the precision and accuracy of the model parameters vary as a function of SNR. To produce data of varying SNR, here the model was fitted to the signal averaged over 1-1000 voxels, producing an SNR range of ~ 16.5 — 520. The fitting was repeated 20 times per SNR. At low SNR, the parameter estimation is highly sensitive to noise and the parameters are degenerate, as indicated by a large spread (standard deviation) in the parameter distributions (see also green box). The precision of the model parameters increases with SNR and the parameter degeneracies are largely overcome when SNR ≳ 100. Although an SNR of ≳ 100 is often not realised *in vivo* (where the *in vivo* data in this study have an SNR of ~ 20), Figure 6 motivates parameter estimation through signal averaging or, as is done for the in vivo data in this study, the concatenation of signal across voxels.

**Figure 6:**
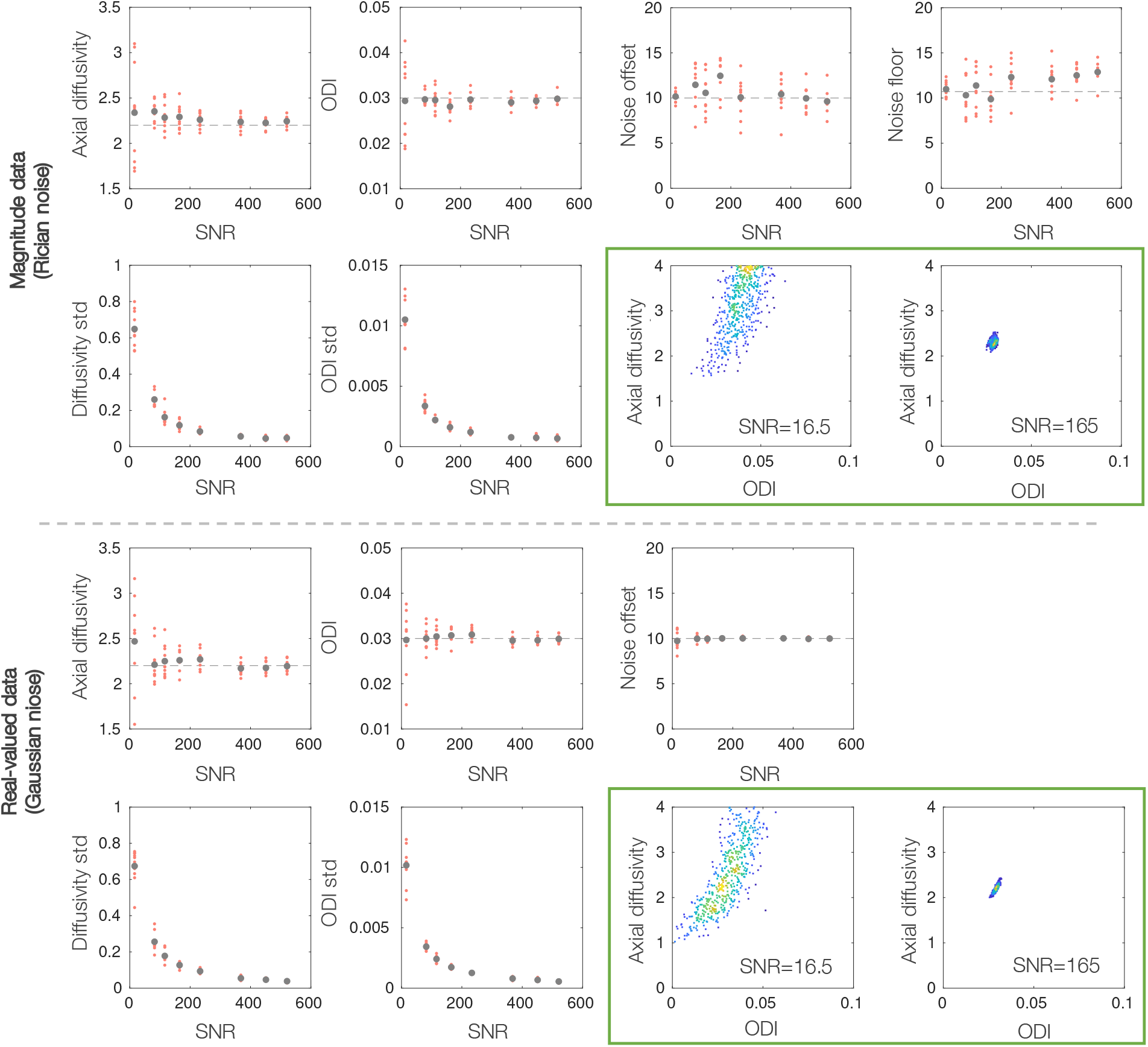
The precision and accuracy of parameter estimates as a function of SNR. The modified NODDI model was fitted to simulated data with a known ground truth (grey dashed line). In high SNR real-valued data, the estimated parameters are estimated more precisely (smaller standard deviation) and tend towards the ground truth values. In magnitude data, the axial diffusivity tends to be overestimated, even at high SNR. The green boxes show the output from MCMC at the SNR of a single voxel and 100 voxels. We see how low SNR leads to parameter degeneracy and biased parameter estimates, both of which are overcome with increased SNR. The diffusivities are given in units of μm^2^/ms.

In magnitude data, the axial diffusivities appear biased and overestimated with respect to the ground truth. This may be related to the calculation of the spherical mean whilst correcting for the rectified noise floor 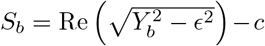, where the positivity constraint can lead to an overestimation of 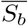. In comparison, parameter accuracy is increased in real-valued data, where the mean axial diffusivity is generally more aligned with the ground truth value.

Together, Figures 5 and 6 motivate the use of real-valued data for more reliable and accurate estimation of axial diffusivity and orientation dispersion via the modified NODDI model.

#### 3.2.3. Assuming incorrect noise characteristics biases parameter estimates

In low SNR regimes, assuming the wrong noise distribution (Rician/Gaussian), or neglecting the presence of a signal offset, could have a considerable effect on the estimated model parameters. Consequently, Figure 7 uses simulated data to examine the effect of getting either of the noise parameters *c* and *ϵ* wrong. We examine both what happens when we neglect the Rician noise floor and signal offset completely (*c, ϵ* = 0), but also what happens when we get them only slightly incorrect. The latter could illustrate a situation where the noise parameters are set to global, predefined values, rather than estimating them on a voxelwise basis to account for local parameter fluctuations. In Figure 7a,b, data were simulated with a signal offset *c* =10 and the model was fitted assuming some fixed offset. In Figure 7c,d, magnitude data were simulated with *c* = 0 and fitted for a fixed noise floor. Note, when the assumed noise floor = 0, this is equivalent to assuming Gaussian noise (Figure 7d). In Figure 7 a,c, even a small error in the noise parameter leads to biased estimates of axial diffusivity (overestimation) and ODI. This is concurrent with increased parameter degeneracy, as indicated by the high standard deviation in the parameter distributions (bottom). In Figure 7c, the ground truth noise floor indicates the standard deviation of the complex Gaussian noise. This value is equivalent to what would be typically measured using de-noising methods and input to Koay inversion method to account for Rician bias. In contrast, the noise floor value which produces an axial diffusivity and ODI closest to the ground truth values is higher. This difference is likely related to Koay’s inversion method being only well suited to data with SNR> 2, where the diffusion signal along many gradients at high b-value will likely not meet this requirement. Again, these results point to potential pitfalls when using Koay’s inversion method to correct for Rician bias using a priori estimates of the noise. In Figure 7b, where *c* is fixed but *ϵ* estimated in magnitude data, a small error in the signal offset c does not have a large effect on the other parameters. This is likely due to the correlation between *c* and *ϵ* (Figure 5), where *ϵ* can be adjusted to account for the error in *c*.

**Figure 7:**
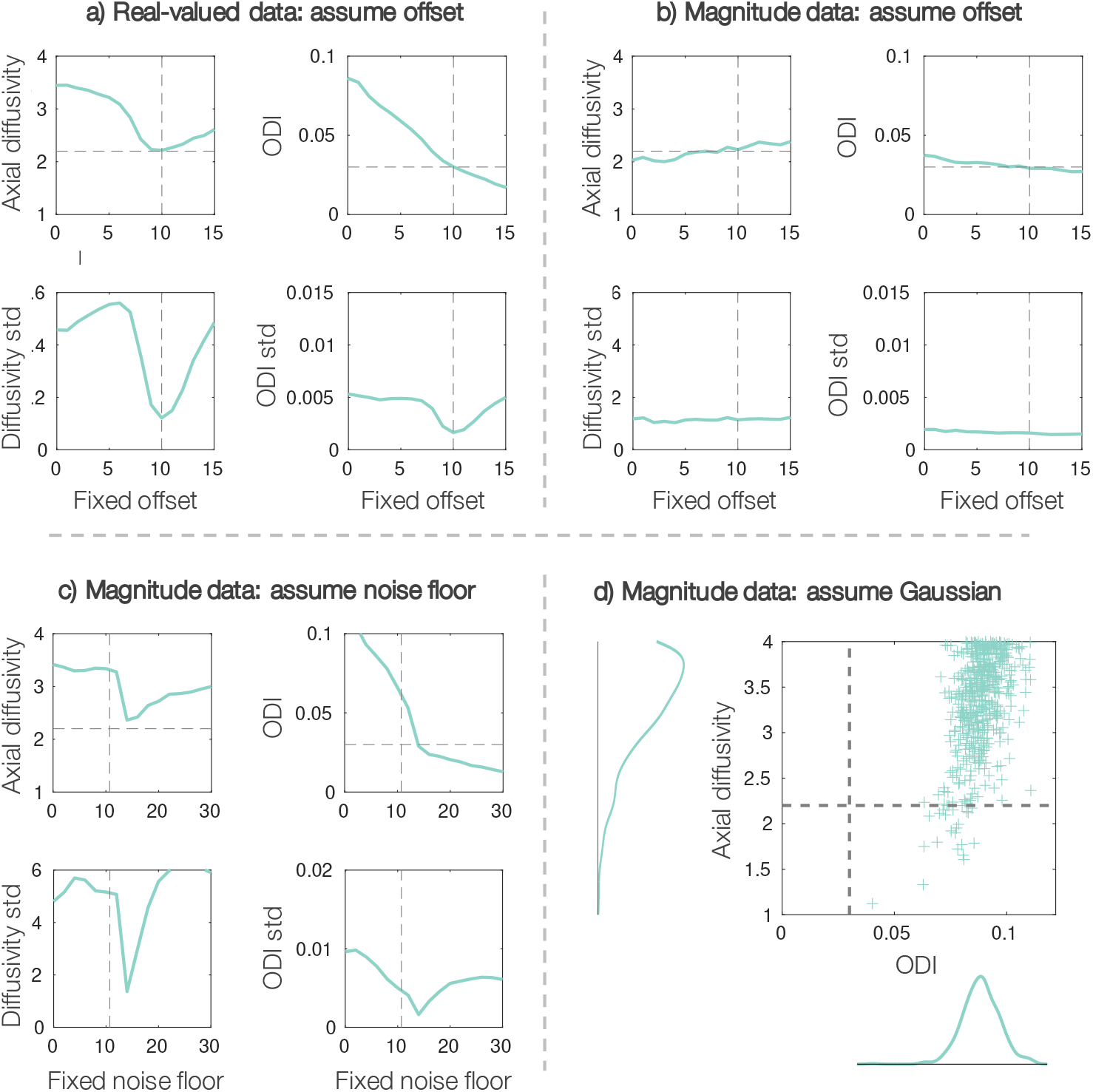
Assuming inaccurate noise parameters biases the estimated axial diffusivity and ODI. Top: data were simulated with *c* =10 and fitted assuming some fixed offset. Bottom: Simulated data with offset *c* = 0 was fitted assuming a fixed noise floor ∈. The plots show the mean (top) and standard deviation (std, bottom) of the parameter distribution output from MCMC. The model was fitted to high SNR data with *N* = 100 and the diffusivities are given in units of μm^2^/ms.

Under the wrong noise model (Figure 7d), the ODI and axial diffusivity estimates are both highly biased (overestimated) and correlated (degenerate), even though the SNR of the data is high (*SNR* ~ 165). When considering the opposite situation i.e. assuming a Rician noise model when fitting to real-valued data, the Rician noise floor e tended to zero as expected, and parameter estimation was otherwise largely unaffected.

#### 3.3. Modified NODDI model for high b-value data: in vivo data

The modified NODDI model was then fitted to high b-value *in vivo* data from 6 healthy subjects. Here we fitted to both magnitude and real-valued images reconstructed from the same complex data. To increase the SNR, data was concatenated across *N* voxels from the corpus callosum, chosen to have similar *S*_0_ and high FA. The results are shown in Figure 8, where the colours indicate data from different subjects.

**Figure 8:**
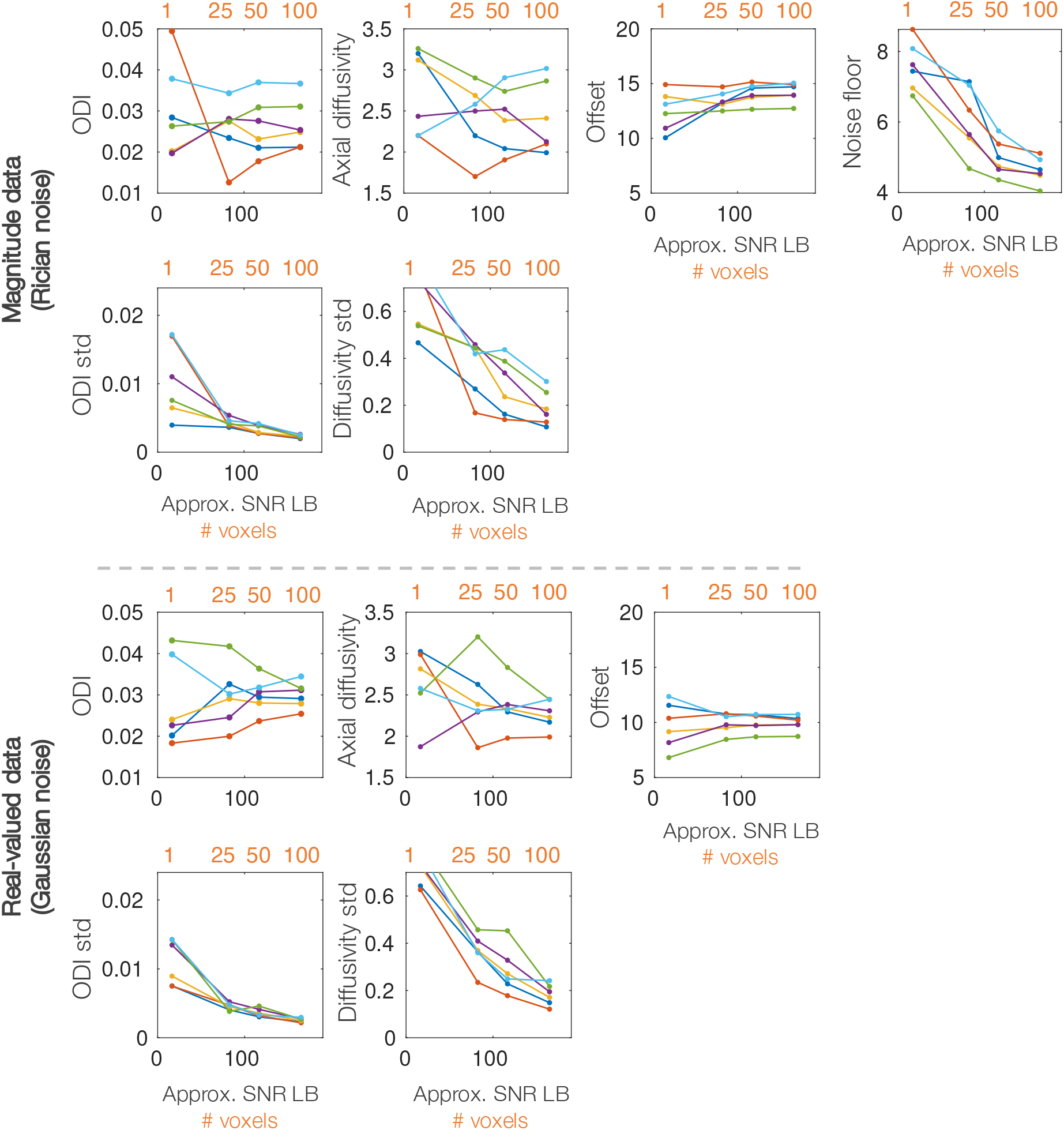
Fitting the modified NODDI model to both magnitude and real-valued *in vivo* data. The model was fitted to the concatenated signal across N=1, 25, 50 or 100 voxels. This corresponds to an approximate SNR given on the x-axis (lower bound, assuming SNR_*vox*_ = 16.5). Plotted are the mean and standard deviation (std) of the parameter distributions output from MCMC. Each colour represents data from a single subject. The diffusivities are given in units of μm^2^/ms.

As in simulated data (Figure 6), the spread of the ODI/diffusivity standard deviation decreases as a function of *N* where in single voxel data the parameters appear degenerate, but can be estimated with increased precision for larger N. Note, whereas in simulated data, each voxel was simulated with identical ODI and axial diffusivity, in *in vivo* data, each voxel is likely to exhibit slightly different diffusivity and fibre orientation distribution. Thus, concatenating signal across many voxels has both the advantage of increasing SNR, and some disadvantage, where the estimated axial diffusivity and ODI are likely some average across the voxel population. The latter was minimised by selecting voxels from within the corpus callosum with similar *S*_0_ and FA. Nonetheless, we generally obtain fairly similar axial diffusivity values when fitting to either N=50 or 100 voxels, giving some confidence to these results.

Both magnitude and real-valued data estimate somewhat similar axial diffusivities and ODI, though the spread of axial diffusivities across subjects is higher for the magnitude data. The mean intra-axonal axial diffusivity across subjects for *N* = 100 was *d*_║_ = 2.26 μm^2^/ms and *d*_║_ = 2.42 μm^2^/ms for real-valued and magnitude data respectively. Crucially, these values are much higher than the *d*_║_ = 1.7 μm^2^/ms typically assumed when fitting NODDI, suggesting that many current studies may be producing biased NODDI parameter estimates.

## 4. Discussion

To describe the complex tissue microstructure in only a handful of modelling parameters, biophysical diffusion models make strong, simplifying assumptions about the underlying tissue architecture. The extent to which many of these assumptions are valid in both healthy and diseased tissue remains an open research question. This study was inspired by the disparity between the estimated intra-axonal axial diffusivity from literature (~ 2 — 2.5 μm^2^/ms [11, 13, 14, 15, 16, 17, 18, 12, 19]) and that typically assumed by the NODDI model and often utilised in simulations (*d*_║_ = 1.7 μm^2^/ms). We first demonstrate the extent to which the NODDI output is dependent on the assumed axial diffusivity. Second, we illustrate how the NODDI framework can be adapted for high b-value data and overcome known parameter degeneracies [2] to estimate axial diffusivity as a free parameter of the model. Crucially, by utilising the NODDI framework and high b-value data, we can forgo modelling of the extra-axonal space and attribute the diffusion attenuation to only two parameters, the intra-axonal axial diffusivity and fibre dispersion, which are simultaneously estimated on a voxelwise basis. Were we to estimate axial diffusivity without accounting for dispersion, our axial diffusivity estimates would be biased [3].

Our results cannot ascertain the “correct” axial diffusivity for the NODDI model, nor do the changes in the NODDI outputs reported here necessarily negate group differences in NODDI parameters. Instead they challenge the interpretation of the NODDI outputs as accurate, biophysical parameters, rather than biased indices which are dependent on many modelling assumptions, including the input axial diffusivity. In Figure 4 – when *d*_║_ = 3 μm^2^/ms (the most extreme case) is compared to *d*_║_ = 1.7 μm^2^/ms – we frequently see the parameter estimates rise or fall by ~ 50%, where in some cases the parameter may be as little as 20% or as much as 250% of its former value. Notably, both the ODI and the signal fractions associated with each of the three compartments change. With higher axial diffusivities, a substantially larger fraction of the white matter signal is associated with the intra-axonal compartment and the associated isotropic compartment is reduced, which may be more in line with our microstructural expectations. This is coupled with an overall ~ 20% increase in the orientation dispersion index, with regions of the corpus callosum and optic tract sometimes doubling. Here, more signal attenuation perpendicular to the fibre is explained by the interplay of intra-axonal diffusion and fibre orientation dispersion, rather than the radial diffusivity associated with the extra-axonal compartment via the tortuosity model.

The change in estimated ODI invites future ODI validation against microscopy gold standards. However, validation is complicated by having to select an appropriate axial diffusivity, a choice that is especially challenging in postmortem tissue where the diffusivities are considerably reduced when compared to their *in vivo* values. In previous work, Schilling *et al.* [49] found the NODDI ODI to correlate well with dispersion values derived from histology (r=0.66), though the ODI was consistently higher than that from histology. In comparison, Grussu *et al.* [1] compared the NODDI outputs to histological data from the human spinal cord to find an approximate one-to-one mapping (r=0.84 in controls, r=0.64 in MS cases). The discrepancy between the results from the two studies may indeed be related to the choice of assumed axial diffusivity. In postmortem spinal cord, Grussu *et al.* assume an axial diffusivity of 1.5 μm^2^/ms which was found to optimise the fitting across all samples. Schilling *et al.* imaged postmortem squirrel monkey brain, though the assumed axial diffusivity wasn’t reported.

Though Figures 3 and 4 assume a single, global diffusivity across the brain and subjects, there is likely great value in being able to account for both between-subject and across-brain variations in intra-axonal axial diffusivity. Recent, promising work by Nilsson et al. [19] utilises data with multiple b-tensor encodings to estimate intra-axonal axial diffusivity on a voxelwise basis, and demonstrate considerable across brain variability, with particularly high axial diffusivities of *d*_║_ ~ 2.7 μm^2^/ms in the corticospinal tract. Studies of intra-axonal axial diffusivity are typically limited to analysing data from relatively few, healthy subjects, making characterisation of normative values or between brain variations challenging. This challenge is likely further exacerbated when considering either development, ageing or pathology which likely either directly or indirectly alter the observed axial diffusivity. Indeed, although the values of axial diffusivity presented here should not be over-interpreted, our results do potentially indicate some across subject variation in intra-axonal axial diffusivity across a series of six healthy participants. Together, these arguments challenge the assumption of a fixed, predefined axial diffusivity and instead advocate for the co-estimation of both architectural features and diffusion characteristics on a voxelwise or subjectwise basis.

A second aim of the study was to demonstrate how by modifying the NODDI model for high b-value data, the framework could be used to simultaneously estimate axial diffusivity and orientation dispersion. Though the NODDI model here explicitly considers only macroscopic fibre orientation, it is likely that microscopic fibre dispersion, including but not limited to axon undulations, also affects intra-axonal axial diffusivity. Indeed Andersson et al. [50] and Lee et al. [51] combine Mote Carlo simulations of diffusion with realistic axon morphologies from 3D X-ray nano-holotomography and electron microscopy data respectively, to demonstrate how complex axon morphology, including axon micro-dispersion, caliber variations, and the presence of mitochondria, all likely influence the observed diffusivities of the tissue. Andersson et al. explicitly show a reduction in the apparent axial diffusivity as a function of the complexity in morphological complexity, with micro-dispersion a key contributing factor.

One of the primary limitations of the NODDI framework is the description of the fibre orientation distribution as that of a single fibre population with symmetric dispersion. Future work should consider modelling multiple fibre populations per voxel and replacing the Watson with a Bingham distribution to account for dispersion asymmetry. Nonetheless, it is reassuring how, even within this restrictive framework, we estimate axial diffusivities from real-valued data in the range *d*_║_ ~ 2 — 2.5μm^2^/ms, with an across subject mean of 2.26μm^2^/ms, in line with current literature [11, 12, 2, 13, 14, 15, 16, 17, 18].

A third aim of this study was to examine the effect of getting the noise characteristics of the data wrong in low SNR data. This is of particular importance as many studies aim to acquire data at high b-value, high spatial resolution, or short scan times – all of which are typically low SNR regimes. When working with our high b-value data, both the presence of a rectified noise floor or signal offset had to be carefully considered to avoid parameter degeneracy and bias. Figure 7 demonstrates the effect of assuming Gaussian noise in magnitude data, as may be often naively done. When the rectified Rician noise floor is not accounted for, the parameter estimates are both biased and correlated (degenerate). This challenges the idea that it is valid to naively apply biophysical models that assume Gaussian noise to low SNR magnitude images. Careful consideration should be taken when transferring models to data with different noise characteristics to what they have been designed and tested on.

Finally, Figure 7 further advocates for estimating noise parameters within the model, rather than as fixed, predefined properties that are e.g. first measured from background voxels and assumed constant across the brain. There are various methods to remove the Rician noise bias that include but are not limited to a) utilising real-valued rather than magnitude images [40, 39], b) approximating the Rician signal as Gaussian via e.g. the Koay inversion technique [37] and, as was not explicitly done in this study, c) applying a Rician noise model during optimisation. Note, the latter requires knowledge of the fibre orientation distribution and so cannot be applied in models fitted to the spherical mean, or powder averaged signal, which explicitly aim to circumvent modelling of the fibre orientation distribution [39]. Further, as the Koay inversion technique is an approximation that breaks down for low SNR data (which likely causes the discrepancy between the ground truth *ϵ* and that estimated from simulated data in Figures 6,7), this study suggests that it may be preferable to use Rician noise modelling rather than Koay’s inversion where possible in low SNR data.

The data used in this study is openly available via http://www.humanconnectomeproject.org/ and [40]. At https://git.fmrib.ox.ac.uk/amyh/noddi-axialdiffusivity we provide a cuDIMOT implementation of NODDI [34] where the assumed diffusivities *d*_║_ and *d_iso_* are user-defined at runtime and MATLAB scripts to implement the modified NODDI model. Interesting avenues for future work include: applying the model to data with both high b-value and high SNR (e.g. from animal models) to explore spatial variations in axial diffusivity; the relationship between axial diffusivity and features of the tissue architecture such as axon diameter; whether intra-axonal axial diffusivity is affected by tissue degradation or disease; the relationship between axial diffusivity and brain development; as well as the estimation of axial diffusivity in postmortem tissue, both in situ and after perfusion or immersion fixation. Furthermore, the model could be extended to estimate multiple fibre populations per voxel and applied brain-wide to provide maps of axial diffusivity across the brain. Finally, co-registered MRI and microscopy data should be used to validate the orientation dispersion estimates reported here. This will likely require the acquisition of a bespoke postmortem dataset which combines data from multiple shells at ultra-high b-values (accounting for the reduced diffusivity of fixed postmortem tissue) with corresponding microscopy imaging of the white matter fibres.

## 5. Conclusion

This study focuses on the assumption of a fixed axial diffusivity in the NODDI model in diffusion MRI. We first demonstrate a considerable dependency of the NODDI parameters on the assumed axial diffusivity, which challenges the interpretation of the NODDI parameters as biophysical metrics, rather than biased indices which are dependent on the modelling assumptions. Second, we demonstrate how the axial diffusivity could be estimated within the NODDI framework by utilising high b-value data. In this challenging, low SNR regime, we demonstrate the importance of locally estimating a rectified noise floor and signal offset i.e. background signal independent of diffusion weighting, both of which could otherwise result in parameter degeneracy or bias. Our results show an axial diffusivity of *d*_║_ ~ 2 — 2.5μm^2^/ms in real-valued *in vivo* human data, which is both in line with current literature and substantially above that typically assumed by NODDI (*d*_║_ = 1.7μm^2^/ms). This motivates the use of more advanced diffusion acquisitions and/or modelling that can resolve parameter degeneracies whilst making minimal assumptions about the tissue microstructure.

## 6. Acknowledgements

AFDH is supported by the EPSRC, MRC and Wellcome Trust (grants EP/L016052/1, MR/L009013/1 and WT202788/Z/16/A). FJL is supported by the Oppenheimer Memorial Trust, St Catherines College Oxford and Wellcome Trust (grant WT215573/Z/19/Z). DKJ is supported by a Wellcome Trust Investigator Award (096646/Z/11/Z) and DKJ and MD by a Wellcome Trust Strategic Award (104943/Z/14/Z). QT is supported by the National Institutes of Health (grant K99-AG073506). KLM and SJ are supported by the Wellcome Trust (grants WT202788/Z/16/A, WT215573/Z/19/Z and WT221933/Z/20/Z). Research reported in this manuscript was supported by the National Institutes of Health under award numbers U01EB026996, P41EB030006, K23NS096056, and R01NS118187. Funding support was also received from the National Natural Science Foundation of China (No.82071994). The Wellcome Centre for Integrative Neuroimaging is supported by core funding from the Wellcome Trust (203139/Z/16/Z).

## 7. Supplementary material

**Supplementary Figure 1:**
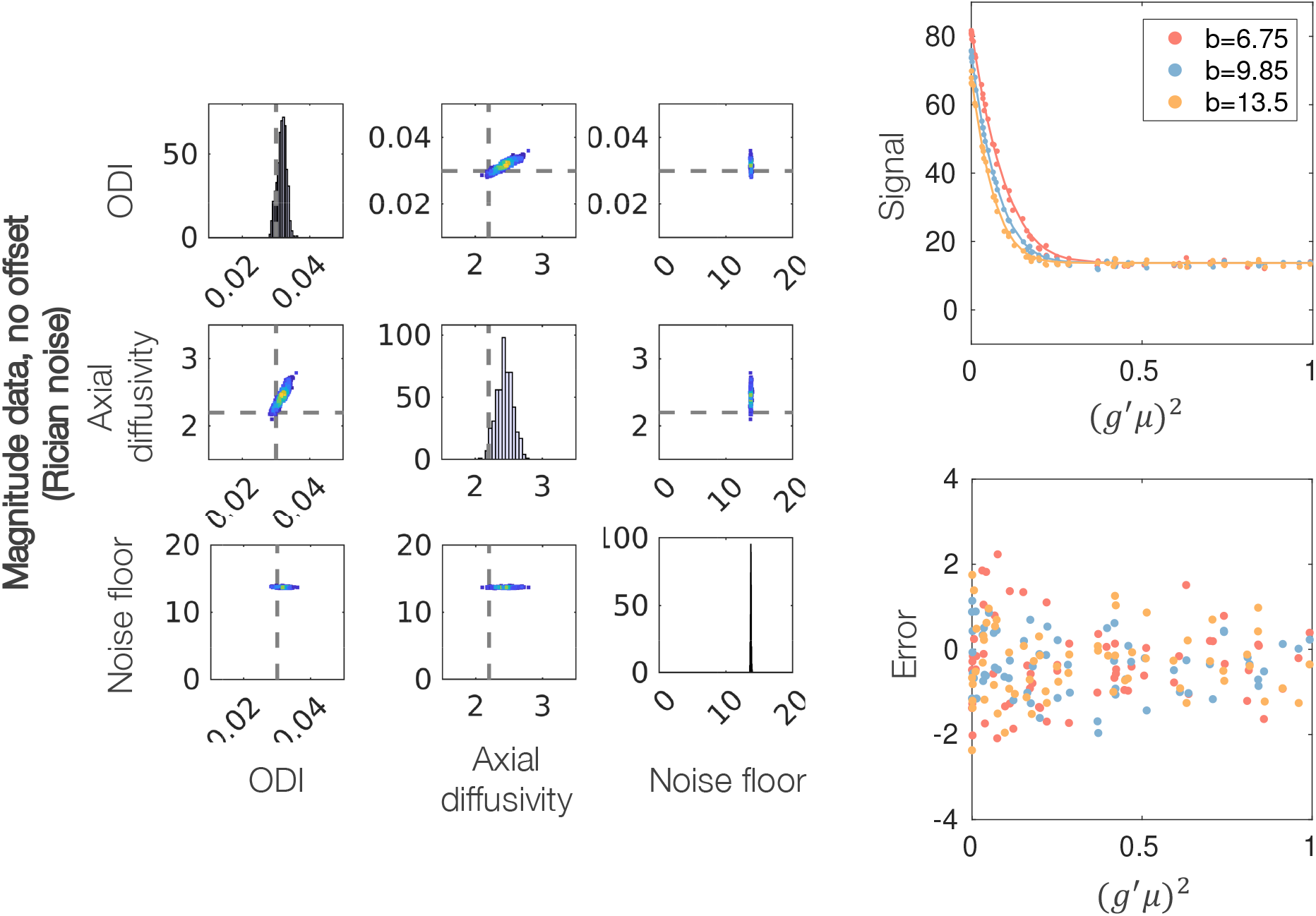
Left: Parameter distributions output from MCMC for magnitude data in a simplified model where the signal offset *c* = 0 and *c* is not estimated as a parameter of the model. Each data point in the scatter plots represents a combination of parameters that fits the signal equally well. Grey dashed lines represent ground truth values. Right: Model fit (line) to simulated data (dots) and the associated error (prediction-data). *g* is the gradient direction, *μ* the fibre orientation and *b* the b-value in ms/μm^2^. The signal is averaged over 100 voxels, each with SNR=16.5.

